# Orally Bioavailable SARS-CoV-2 Protease Inhibitors Bearing a Hydroxymethyl Ketone Warhead

**DOI:** 10.64898/2026.05.15.725542

**Authors:** N. G. R. Dayan Elshan, Karen C. Wolff, Frank Weiss, Sourav Ghorai, Gennadii Grabovyi, Katy Wilson, Laura Riva, Ashley K. Woods, James Pedroarena, Armen Nazarian, Yuyin Liu, Wrickban Mazumdar, Lirui Song, Neechi Okwor, Jacqueline Malvin, Malina A. Bakowski, Melanie G. Kirkpatrick, Amal Gebara-Lamb, Edward Huang, Vân T. B. Nguyen-Tran, Victor Chi, Shuangwei Li, Kyoung-Jin Lee, Case W. McNamara, Anil Kumar Gupta, Alireza Rahimi, Jian Jeffrey Chen, Sean B. Joseph, Peter G. Schultz, Arnab K. Chatterjee

## Abstract

The use of covalent warheads targeting the catalytic cysteine has been a cornerstone in coronavirus main protease (M^pro^) inhibitor development, where various electrophilic motifs have been used including aldehydes, nitriles, ketoamides, and hydroxymethyl ketones (HMKs). Recent efforts have been mostly centered around nitrile warheads, given the success of compounds like Nirmatrelvir and Ensitrelvir in the clinic. However, finding and advancing alternative chemotypes with differentiating chemical and pharmacological profiles is essential for future pandemic preparedness. Among such alternatives, HMKs hold special interest because they balance reduced intrinsic electrophilicity with an excellent selectivity profile. Nevertheless, early HMK-based compounds, such as the clinical-stage M^pro^ inhibitor PF-00835231, suffered from poor oral bioavailability and therefore required intravenous administration, with or without prodrug derivatization of the hydroxyl group. Here, we describe our efforts in advancing the HMK field via the discovery of **mCMX110**, a lead that has superior potency, increased unbound exposure *in vivo*, and favorable oral bioavailability in preclinical studies.

**Graphical Abstract:** 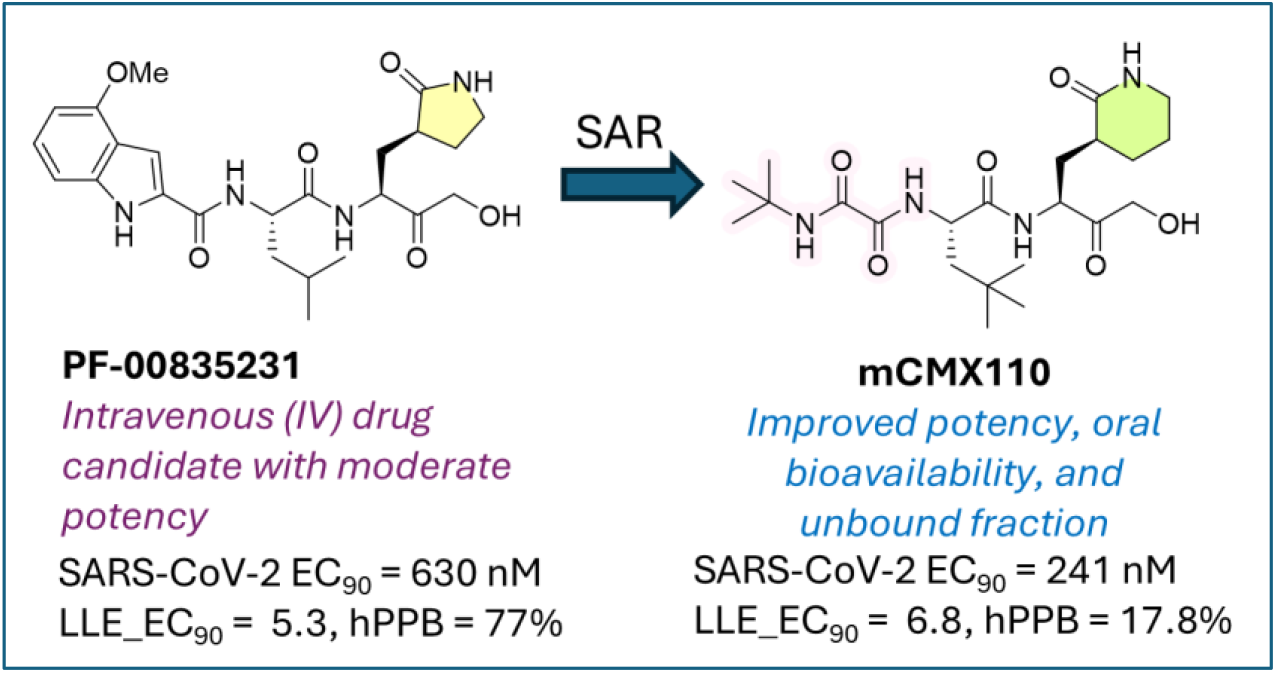

## 1. INTRODUCTION

Severe acute respiratory syndrome coronavirus 2 (SARS-CoV-2) created significant disruption of everyday life and led to substantial mortality across the globe. Efforts by researchers over the last several years have resulted in vaccines and antiviral agents that have significantly reduced morbidity and mortality rates from SARS-CoV-2. Despite this, the fundamental risks of viral pandemics remain a global health concern among the scientific community due to continued viral evolution and the risk of emerging variants with altered transmissibility and therapeutic susceptibility. The development of next-generation antivirals targeting conserved viral proteins is one of the most attractive ways to address this risk, ensuring preparedness against current and future coronaviruses.

The SARS-CoV-2 main protease (M^pro^, also termed 3CL^pro^) is an essential cysteine protease responsible for proteolytic processing of viral polyproteins (pp1a and pp1ab) into functional nonstructural proteins required for replication and transcription.^1^ M^pro^ cleaves at multiple conserved Leu-Gln↓(Ser,Ala,Gly) motifs and lacks closely related human homologs, making it an attractive antiviral target with reduced risk of off-target toxicity.^1^ Structural studies have revealed a catalytic Cys145–His41 dyad within a well-defined substrate binding pocket, enabling rational design of covalent and reversible covalent inhibitors.^1-3^

Covalent warheads targeting the catalytic cysteine have been particularly successful in M^pro^ inhibitor development, particularly exemplified by Nirmatrelvir and Ensitrelvir. Among these, electrophilic motifs including aldehydes, nitriles, ketoamides, and hydroxymethyl ketones (HMKs) form reversible covalent adducts with Cys145, stabilizing a tetrahedral hemithioacetal intermediate that mimics the transition state of peptide cleavage.^4^ Hydroxymethyl ketones are of special interest because they balance reactivity and selectivity: they are sufficiently electrophilic to engage catalytic cysteines while typically exhibiting improved metabolic stability and reduced nonspecific reactivity compared to other warheads.^4^ HMK warheads have been previously validated versus cysteine proteases, supporting their suitability for targeting coronavirus proteases.^4-6^ PF-00835231, the first-in-class SARS-CoV-1 Mpro inhibitor developed by Pfizer is one of the key examples of a HMK-based drug candidate, but lacked the oral bioavailability to cover efficacious exposure against M^pro^. At the onset of the SARS-CoV-2 pandemic, PF-00835231 (**Figure 1**) was modified into a prodrug form (PF-07304814, Lufotrelvir, **Figure 1**)^6^ and evaluated as an intravenous treatment in the clinic. Further development of these HMK-based M^pro^ inhibitors was abandoned with the discovery of orally bioavailable M^pro^ inhibitors such as Nirmatrelvir.

**Figure 1:**
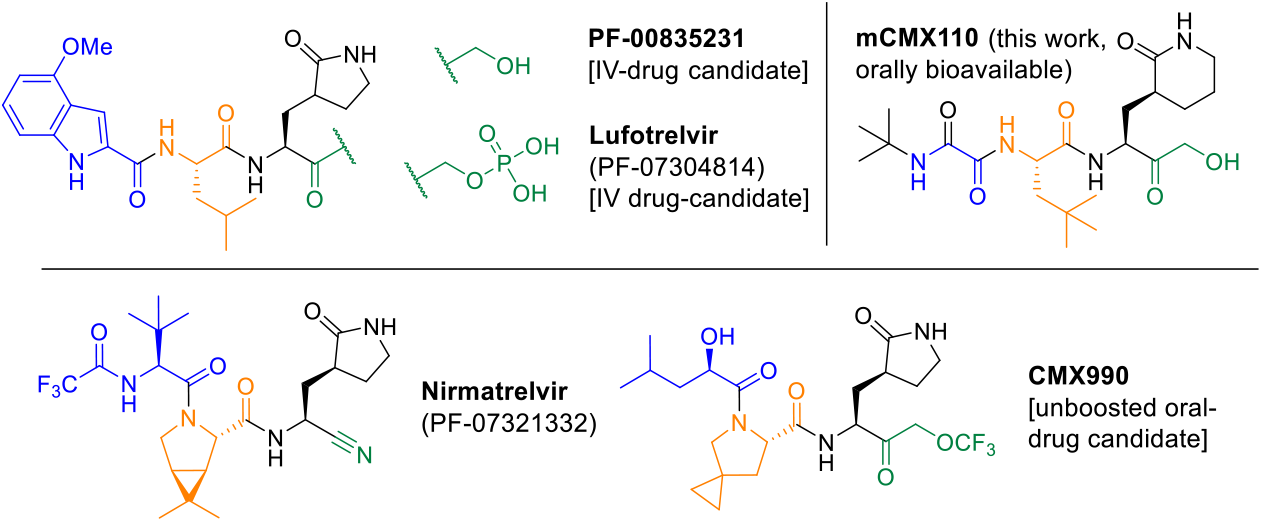
Synthesis of mCMX110. (A) synthesis of HMK warhead containing P1-motif. (B) final assembly of **mCMX110** (19% overall yield). This scheme is representative of the synthesis of all HMK compounds described in this text; and has been used to synthesize material in gram scale.

At present, the clinical landscape of SARS-CoV-2 M^pro^ inhibitors is dominated by strong covalently associating warheads such as nitriles or emerging warheads as seen in our recent clinical compound CMX990^7^. These work well in many circumstances, but still have caveats to their overall pharmacokinetics, efficacy, and safety profile (e.g., Nirmatrelvir requires co-dosing with CYP-inhibitor ritonavir to boost exposure for target coverage). These shortcomings, the risks of resistance driving mutations emerging in M^pro^,^8, 9^ and in an overall view of preparedness against future pandemics, it is essential that we discover novel, structurally diverse chemotypes to expand the chemical space of M^pro^ inhibitors. To this end we disclose here another chemical series in our SARS-CoV-2 M^pro^ inhibitor program, outlining the discovery of **mCMX110**, an HMK warhead bearing M^pro^ inhibitor that differentiates from PF-00835231 in having superior oral bioavailability without a need for ritonavir-boosting or prodrug strategies.

## 2. Materials and methods

All reagents and solvents were purchased from commercial sources and used without further purification unless otherwise specified. Flash column chromatography was performed using silica gel (200–300 mesh). All reactions were monitored by TLC (pre-coated EMD silica gel 60 F254 TLC aluminum sheets and visualized with a UV lamp or appropriate stains) and/or LCMS (Waters Acquity UPLC system, 2 or 4 min run of a 10–90% mobile phase gradient of acetonitrile in water [+0.1% formic acid]). NMR spectra were obtained on Bruker AV400 or AV500 instruments, and data was analyzed using the MestReNova NMR software (Mestrelab Research S. L.). Chemical shifts (δ) are expressed in ppm and are internally referenced for ^1^H NMR (DMSO-*d*_*6*_ 2.50 ppm) and ^13^C NMR (CDCl_3_ 77.16 ppm, DMSO-*d*_*6*_ 39.52 ppm). X-ray data was collected at room temperature on a Bruker D8 QUEST instrument with an IμS Mo microfocus source (λ = 0.7107 Å) and a PHOTON-III detector.

### Chemistry

All compounds with the hydroxymethylketone warhead were synthesized according to the representative procedure outlined for **mCMX110** (**Scheme 1**). All compounds described are >95% pure by HPLC and/or NMR analyses.

#### ((*S*)-2-((*tert*-butoxycarbonyl)amino)-3-((*S*)-2-oxopiperidin-3-yl)propanoic acid (2)

To a stirred solution of methyl (*S*)-2-((tert-butoxycarbonyl)amino)-3-((*S*)-2-oxopyrrolidin-3-yl)propanoate 1 (10 g, 33.3 mmol, 1.0 eq) in tetrahydrofuran (70 mL), lithium hydroxide monohydrate (1.68 g, 40.0 mmol, 1.2 eq) in water (30 mL) was added at 0 °C and the mixture was stirred 1 h. After completion, the reaction mixture was cooled 0 °C, acidified with 2 N hydrochloric acid (pH∼5) and extracted with ethyl acetate (3 × 300 mL). The combined organic layer was dried over anhydrous sodium sulphate and concentrated under reduced pressure to afford the free acid **2** (8.0 g, 30.0 mmol, 90%) as a pale yellow gum. TLC system: MeOH:DCM (1:9); *R*_*f*_ : 0.1.

#### *tert*-butyl ((*S*)-4-bromo-3-oxo-1-((*S*)-2-oxopiperidin-3-yl)butan-2-yl)carbamate (3)

To a stirred solution of **2** (8.0 g, 30.0 mmol, 1.0 eq) in tetrahydrofuran (100 mL), *N*-methylmorpholine (6.58 mL, 60.0 mmol, 2.0 eq) was added then isobutyl chloroformate (6.13 mL, 44.9 mmol, 2.2 eq) was added at -10 °C and the reaction mixture was stirred at -10 °C for 1 h. After completion, the reaction mixture was filtered and washed with tetrahydrofuran (50 mL). Freshly prepared diazomethane in diethyl ether (prepared from 4.0 eq. of diazald) was added to the filtrate at -10 °C and the reaction mixture was stirred at room temperature for 1 h. After completion, the reaction mixture was diluted with water (100 mL) and extracted with ethyl acetate (3 × 300 mL). The combined organic layer was dried over anhydrous sodium sulphate and evaporated under reduced pressure to afford *tert*-butyl ((*S*)-4-diazo-3-oxo-1-((*S*)-2-oxopiperidin-3-yl)butan-2-yl)carbamate (9.8 g, crude) as a yellow liquid. TLC system: EtOAc : pet-ether (7:3); *R*_*f*_ : 0.2.

To a stirred solution of the α-diazo derivative above (9.8 g, 31.6 mmol) in tetrahydrofuran (100 mL) was added 48% aqueous hydrobromic acid (6.4 mL, 37.94 mmol) dropwise at -10 °C and the mixture was stirred at -10 °C for 30 min. After completion, the reaction mixture was basified with saturated sodium bicarbonate and extracted with ethyl acetate (3 × 150 mL). The combined organic layer was dried over anhydrous sodium sulphate and concentrated under reduced pressure. The crude residue was then purified by column chromatography over silica gel (230-400 mesh) using 50-70% ethyl acetate in pet-ether as a gradient to afford *the* α-bromomethyl-ketone derivative **3** (5.4 g, 14.9 mmol, 50% over two steps) as a yellow liquid. TLC system: EtOAc : pet-ether (7:3); *R*_*f*_ : 0.4.

#### *tert*-butyl ((*S*)-4-hydroxy-3-oxo-1-((*S*)-2-oxopiperidin-3-yl)butan-2-yl)carbamate (**4**)

To the stirred solution of **3** (4.8 g, 14.9 mmol, 1.0 eq) in THF:H_2_O (35:15 mL), sodium bicarbonate (6.26 g, 74.6 mmol, 5.0 eq) was added and the reaction mixture was stirred at room temperature for 24 h. After completion, the reaction mixture was diluted with water (100 mL) and extracted with ethyl acetate (3 × 150 mL). The combined organic layer was dried over anhydrous sodium sulfate and evaporated under reduced pressure to give crude product. The crude residue was then purified by silica gel (230-400 mesh) column chromatography using 5-7% MeOH in DCM as a gradient to afford α-hydroxymethyl-ketone **4** (2.3 g, 7.7 mmol, 51%) as an off-white solid. TLC system: EtOAc:pet-ether (7:3); *R*_*f*_: 0.5.

#### (*S*)-3-((*S*)-2-amino-4-hydroxy-3-oxobutyl)piperidin-2-one hydrochloride (**5**)

A solution of **4** (3 g, 10.0 mmol, 1 eq) in DCM (45 mL) was treated with HCl/dioxane(4N) (10 mL, 40.0 mmol, 4.0 eq) for 15 min at 0 °C under nitrogen. The resulting mixture was stirred for 1 h at room temperature under nitrogen atmosphere. After the reaction was completed, the solvent was removed under vacuum to give the hydrochloride salt **5** (2.1 g, quant.) as a white solid. LCMS:(ES,m/z): [M+H]^+^=201.

#### methyl 2-(*tert*-butylamino)-2-oxoacetate (**6**)

To a stirred mixture of tert-butylamine (75 g, 1025 mmol, 1.0 eq) and TEA (311.30 g, 3076.3 mmol, 3 eq) in DCM (750 mL) was added methyl oxalochloridate (150.74 g, 1230.5 mmol, 1.2 eq) dropwise at 0 °C under nitrogen atmosphere. The resulting mixture was washed with 1N HCl (5×500 mL), brine (2×200 mL), dried over anhydrous Na_2_SO_4_. After filtration, the filtrate was concentrated under reduced pressure. The residue was purified by silica gel column chromatography, eluted with n-hexane/EA (1:1) to afford methyl ester **6** (130 g, 902 mmol, 88%) as a yellow oil. LCMS:(ES,m/z): [M+H]^+^=160.

#### (2*S*)-2-[(*tert*-butylcarbamoyl) formamido]-4,4-dimethylpentanoic acid (7)

To a stirred solution of 4-methyl-L-leucine (2.9 g, 20.0 mmol, 1.0 eq) and **6** (3.82 g, 24.0 mmol, 1.2 eq) in ACN (70 mL) were added DBU (7.60 g, 49.9 mmol, 2.5 eq) at 0 °C under nitrogen atmosphere. The resulting mixture was stirred for 1 h at room temperature under nitrogen atmosphere. After the reaction was completed, the mixture was diluted with EtOAc (200 mL), washed with 1N HCl (5×70 mL), brine (3 × 70 mL) and dried over anhydrous Na_2_SO_4_. After filtration, the filtrate was concentrated under reduced pressure. The residue was purified by silica gel column chromatography, eluted with PE/THF (1:1) to afford the acid **7** (3.2 g, 11.7 mmol, 59%) as a brown yellow solid. LCMS:(ES,m/z): [M+H]^+^=273. ^1^H NMR (300 MHz, DMSO-*d*_*6*_): *δ* 8.75-8.72 (m, 1H), 7.82(s, 1H), 4.27-4.21(m,1H), 1.87-1.68(m, 2H), 1.33(s, 9H), 0.88(s, 9H).

#### *N*-*tert*-butyl-*N*’-[(1S)-1-[[(2*S*)-4-hydroxy-3-oxo-1-[(3*S*)-2-oxopiperidin-3-yl] butan-2-yl] carbamoyl] -3,3-dimethylbutyl] ethanediamide (mCMX110)

To a stirred solution of **7** (2.1 g, 7.7 mmol, 1.0 eq) and the hydrochloride **5** (2.19 g, 9.3 mmol, 1.2 eq) in DCM (70 mL) were added HATU (3.52 g, 9.2 mmol, 1.2 eq) and DIEA (2.49 g, 19.3 mmol, 2.5 eq) at 0 °C under nitrogen atmosphere. The resulting mixture was stirred for 1 h at room temperature under nitrogen atmosphere. After the reaction was completed, it was quenched with 10 mL water at 0 °C. The resulting mixture was washed with 1N HCl (5x70mL). The combined organic layers were washed with brine (50 mL), dried over anhydrous Na_2_SO_4_. After filtration, the filtrate was concentrated under reduced pressure. The residue was purified by silica gel column chromatography, eluted with PE/THF (1:3) to afford **mCMX110** (1.3 g, 2.9 mmol, 37%) as a white solid. It was further purified by SFC to give 0.7 g the final target as white amorphous solid.

#### SFC separation condition

Column: CHIRALPAK IH, 3*25 cm, 5 μm; Mobile Phase A: CO2, Mobile Phase B: IPA: ACN=1: 1; Flow rate: 80 mL/min; Gradient: isocratic 30% B; Column Temperature (°C): 35; Back Pressure(bar): 100; Wave Length: 220 nm; RT1(min): 2.63; RT2(min): 5.27; Sample Solvent: IPA; Injection Volume: 5 mL; Number Of Runs: 10)

^6^HRMS m/z *calcd*. for C_22_H_39_N_4_O_6_ ^+^ [M + H]^+^ 455.2864, found 455.4780; ^1^H NMR (400 MHz, DMSO-*d*_*6*_): *δ* 8.61 (d, *J* = 9.0 Hz, 1H), 8.39 (d, *J* = 8.0 Hz, 1H), 7.84 (s, 1H), 7.46 (s, 1H), 5.09 (t, *J* = 6.0 Hz, 1H), 4.51 – 4.41 (m, 1H), 4.38 – 4.29 (m, 1H), 4.26 – 4.04 (m, 2H), 3.15 – 3.05 (m, 2H), 2.18 – 2.03 (m, 2H), 1.87 – 1.45 (m, 6H), 1.39 – 1.21 (m, 10H), 0.88 (s, 9H).

### *In vitro* Biology

The following viral strains were obtained through BEI Resources, NIAID, NIH: Isolate HCOV-19 USA-WA1/2020 (NR-52281), Isolate hCoV-19/USA/OR-OHSU-PHL00037/2021 (Alpha Variant, NR-55461), Isolate HCOV-19/USA/PHC658/2021 (Delta Variant, NR-55691), isolate HCOV-19/USA/MD-HP20874/2021 (Omicron Variant, NR-56461), Human Coronavirus, OC43 (NR-52725), and Human Coronavirus, 229E (NR-52726). The SARS-CoV-2 assays (cell-based, enzymatic) were conducted as described previously.^7, 10^ HCoV-OC43 was propagated in HCT-8 cells (ATCC CCL-244) and compound activity was assayed in a high-content imaging assay after a 48 hr incubation using the mouse monoclonal antibody OC-43 strain clone 541-8F (Sigma Millipore MAB9012) and goat anti-mouse H+L conjugated Alexa 488 secondary (Thermo Fisher Scientific A11001). HCoV-229E was propagated in MRC-5 pd25 cells (Sigma cat # 05081101-1VL) and compound activity determined by assessing cell viability at 120 hr post-infection using CellTiter-Glo® (Promega No G7573). The uninfected cytotoxicity counter screens were incubated in parallel with the antiviral assays and cell viability assessed using CellTiter-Glo®.

### Drug Metabolism and Pharmacokinetics (DMPK) Assays

Described *in vitro* ADME assays were conducted at Calibr-Skaggs (San Diego, CA) or at WuXi AppTec (Shanghai, China) using the procedures previously described.^7, 10^

Animal experimental procedures were approved by the Institutional Animal Care and Use Committee (IACUC) of the respective study locations. Pharmacokinetic studies were conducted at Aragen (India), Calibr (San Diego, CA), WuXi (China), or Pharmaron Inc. (China)

Pharmacokinetics in animals [CD1-mouse, SD-rat, Syrian hamster, beagle dog]:

For all reported intravenous (IV) pharmacokinetic studies in this manuscript, three fasted animals per study group were administered the test article as a solution in 50% PEG 300, 10% ethanol, and 40% saline. For the oral PK studies, same formulation was used for solutions; whereas for suspensions – 0.5% methylcellulose (MC) or 0.5% Sodium carboxymethyl cellulose (NaCMC) in water was used. Blood samples were collected per general protocol at 0.083, 0.5, 1, 3, 5, 8, and 24 hr post-dosing for IV studies; and at 0.5, 1, 3, 5, 8, and 24 hr post-dosing for PO studies.

The blood samples were centrifuged to obtain the plasma, which was stored below -20°C until analysis. Plasma concentrations were determined by liquid chromatography/tandem mass spectrometry (LC−MS/MS). The PK parameters were determined by non-compartmental methods using WinNonLin (v6.1 or higher version, Certara Inc.).

## 3. Results

### Syntheses of mCMX110 series is readily feasible

Synthesis of this hydroxymethyl ketone compound series follows generally known synthetic chemistry methods described by us and others in literature.^3, 5, 7^ Representative synthesis of **mCMX110** (**Scheme 1**) was achieved in 19% yield over the three steps from *t*-butylamine. We have used the currently disclosed sequence to make **mCMX110** in >20 g scales. Without the use of SFC at the final step, major impurity is the epimerization-derived diastereomer impurity at P1. Based on our prior experiences in similar compounds, this process can be further optimized to avoid the SFC separation needs and improve the overall yield.

**Scheme 1:**
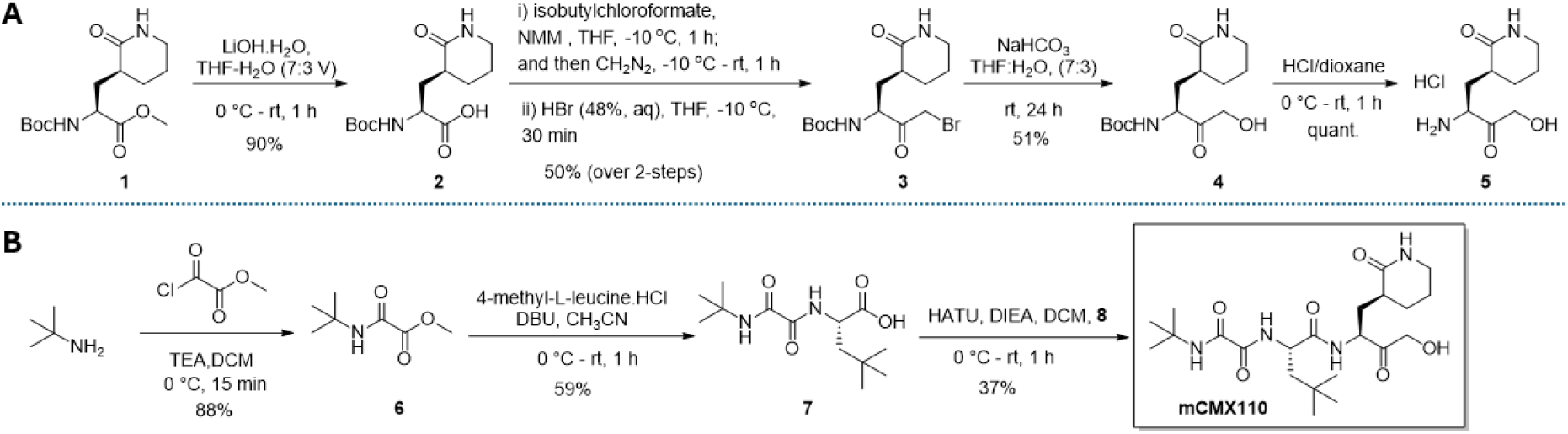
Representative known coronavirus protease inhibitors with various covalent warheads in comparison to the lead mCMX110 from the current work. Generally, up to four distinct structural regions are found in all peptidomimetic CL^pro^ inhibitors: warhead (WH), P1 (amino acid 1, recognition element with a lactam side-chain), P2 (AA 2), and P3 (a *N*-Cap, or a third AA). The warheads can have distinct differences in electrophilicity that contributes substantially to the potency and pharmacokinetic performance of the compound. **mCMX110** (lead from this work) improves upon PF-00835231 by having superior oral bioavailability without a need for ritonavir-boosting or prodrug strategies.

### SAR studies establish mCMX110 to have the best balance between potency and early ADME

**properties:** We synthesized ∼100 analogs in this HMK series, where the key SAR improvements were found through oxamide P2-caps (**Figure 2**). Maintaining the γ-lactam (pyrrolidinone) P1 motif of PF-00835231, and bringing in the aryl oxamide P2-cap in place of the methoxyindole, gave an immediate improvement of potency (>2× improved) and lipophilic ligand efficiency (LLE, 1.4 units improvement) as represented by the compound **8**.

**Figure 2:**
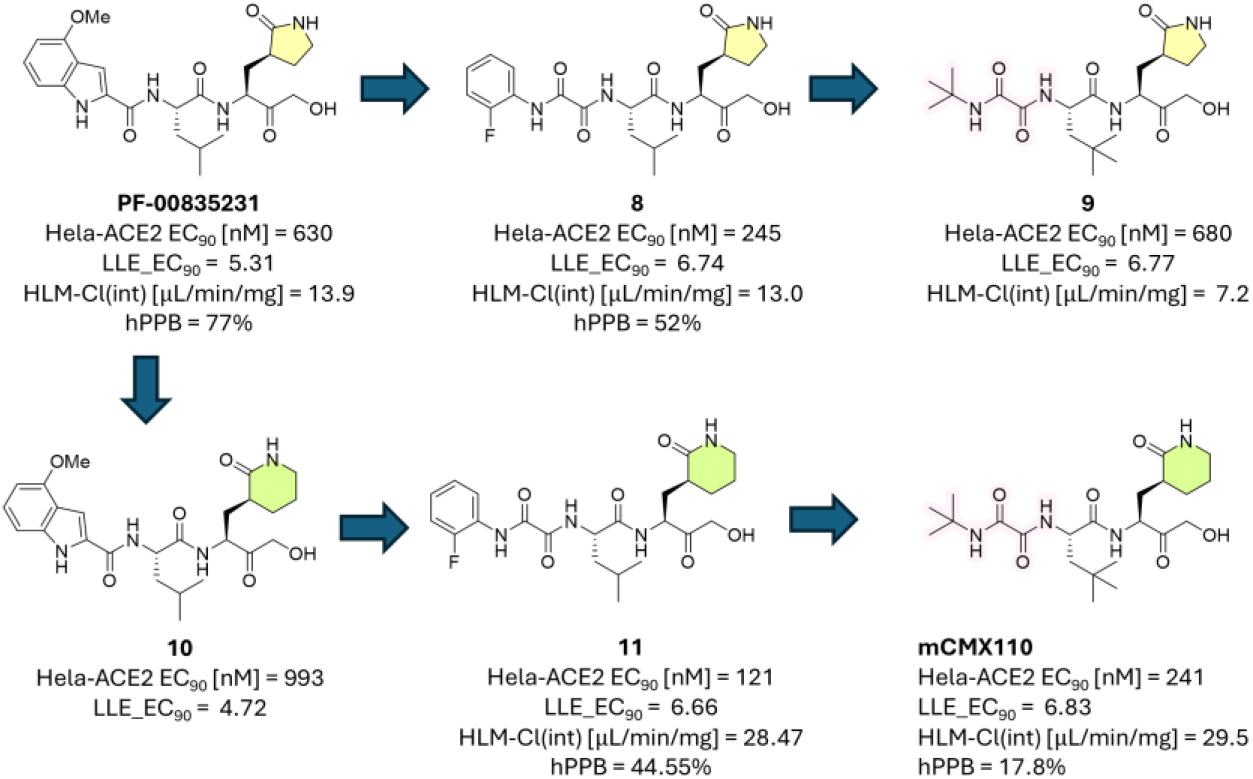
Representative SAR summary trajectory to early tool compound 8, and ultimately to lead mCMX110. SAR across all regions of the compounds were carried out in parallel, leading to a series of ∼100 HMK compounds.

Several aryl-oxamide containing compounds demonstrated positive mini-Ames assay results, upon which we decided to move away from the anilinic oxamides to minimize developmental risk. Among other modifications (**Table 1**), replacement of the para-fluorophenyl group of **8** with a t-butyl group resulted in analog **9** with 2x improvement in microsomal clearance, but compromised potency. To gain the potency back, we envisioned replacement of the pyrrolidinone motif with a piperidinone P1 side-chain. Indeed, in most cases (e.g., **11** vs. **8**), the δ-lactam was superior to the γ-lactam counterpart in driving potency, albeit at a moderate cost of microsomal clearance. Eventually, our SAR exploration exemplified in **Table 1** led to the identification of lead compound **mCMX110**, which retained the potency of **8** without compromising LLE. Importantly **mCMX110** had low plasma protein binding (human PPB almost 3x lower vs **8**), which favorably increased the unbound fraction that is recognized as a key determinant of antiviral efficacy.

**Table 1:**
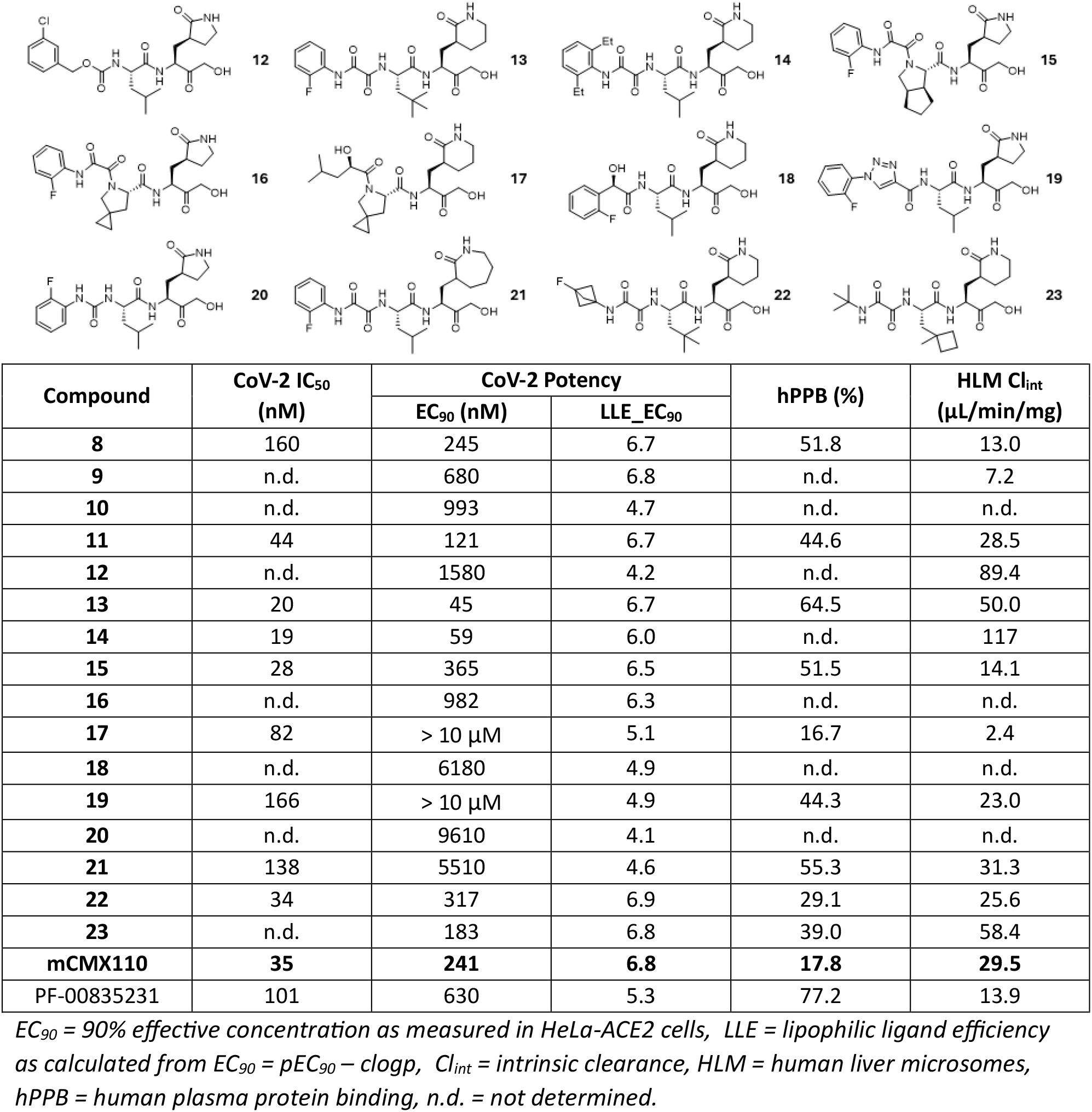
Representative compounds and profiling data for HMK series SAR.

|  |  |  |  |  |  |
| --- | --- | --- | --- | --- | --- |
| Compound | CoV-2 IC<sub>50</sub><br>(nM) | CoV-2 Potency | | hPPB (%) | HLM Cl<sub>int</sub><br>(μL/min/mg) |
|  |  | EC<sub>90</sub> (nM) | LLE_EC<sub>90</sub> |  |  |
| **8** | 160 | 245 | 6.7 | 51.8 | 13.0 |
| **9** | n.d. | 680 | 6.8 | n.d. | 7.2 |
| **10** | n.d. | 993 | 4.7 | n.d. | n.d. |
| **11** | 44 | 121 | 6.7 | 44.6 | 28.5 |
| **12** | n.d. | 1580 | 4.2 | n.d. | 89.4 |
| **13** | 20 | 45 | 6.7 | 64.5 | 50.0 |
| **14** | 19 | 59 | 6.0 | n.d. | 117 |
| **15** | 28 | 365 | 6.5 | 51.5 | 14.1 |
| **16** | n.d. | 982 | 6.3 | n.d. | n.d. |
| **17** | 82 | > 10 μM | 5.1 | 16.7 | 2.4 |
| **18** | n.d. | 6180 | 4.9 | n.d. | n.d. |
| **19** | 166 | > 10 μM | 4.9 | 44.3 | 23.0 |
| **20** | n.d. | 9610 | 4.1 | n.d. | n.d. |
| **21** | 138 | 5510 | 4.6 | 55.3 | 31.3 |
| **22** | 34 | 317 | 6.9 | 29.1 | 25.6 |
| **23** | n.d. | 183 | 6.8 | 39.0 | 58.4 |
| **mCMX110** | **35** | **241** | **6.8** | **17.8** | **29.5** |
| PF-00835231 | 101 | 630 | 5.3 | 77.2 | 13.9 |

EC<sub>90</sub> = 90% effective concentration as measured in HeLa-ACE2 cells, LLE = lipophilic ligand efficiency as calculated from EC<sub>90</sub> = pEC<sub>90</sub> – clogp, Cl<sub>int</sub> = intrinsic clearance, HLM = human liver microsomes, hPPB = human plasma protein binding, n.d. = not determined.

Incorporation of other groups at P2-cap (e.g., carbamate cap analog **12**) most often led to reduced potency. Aniline based aryl-oxamides had by far the best potency improvements, where compounds such as **13** (CoV-2 EC_90_ = 45 nM) managed to push the potencies >14x improved over PF-00835231. Aryl oxamide P2-caps did not pair well with proline P2-derivatives as exemplified by **15** (CoV-2 EC_90_ = 365 nM) and **16** (CoV-2 EC_90_ = 982 nM). They were still better in pairing to P2-prolines versus alternative P3-cap such as the leucic acid P2-cap in **17** (CoV-2 EC_90_ > 10 µM).

In addition to the Ames-liability, aryl-oxamides unfortunately also had general instability in rodent plasma, which was another factor (as rodent toxicology and efficacy studies would have met limiting exposures) in the decision to move away from these potent compounds. We tried to explore isosteric derivatives such as **18** (CoV-2 EC_90_ > 6 µM) or **19** (CoV-2 EC_90_ > 10 µM); with significant losses of potency evident. Removal of one carbonyl motif while retaining similar count of H-bond donors in **20** (CoV-2 EC_90_ > 9 µM) lost significant activity.

Notably, the addition of the extra methyl group at the P2 position of **13** gives a potency advantage of ∼3x over **11**, consistent with the hydrophobic nature of the SARS-CoV-2 Mpro S2 pocket. This, along with the shift to the 6-membered lactam at P1 side-chain, helped **mCMX110** retain the potency of **8** despite the move away from aryl oxamides. Moving to a seven-membered lactam P1 in **21** (CoV-2 EC_90_ > 5 µM) did not improve potency. We attempted to incorporate more constrained structural features to mCMX110 as highlighted by bicyclo[1.1.1]butane derivative **22** and cyclobutyl derivative **23**. Such derivatizations failed to provide a profile superior to **mCMX110** on the balance between potency, microsomal clearance, and unbound fraction.

### Formulation, stability, and solubility profile of mCMX110 to enable oral administration

In general,most PK studies described in this manuscript for **mCMX110** were run with the amorphous material (DSC T_g_ onset ∼66 °C) obtained post SFC purification. **mCMX110** however can be crystallized as a stable single polymorph (DSC melting onset = 86.3 °C, peak temperature =91.6 °C), using an antisolvent crystallization approach (acetone/pentane). Bulk stability of this crystallized material is moderate, where **mCMX110** demonstrates good stability under humid conditions at 40 °C over short durations, but undergoes substantial degradation at elevated temperature (60 °C) and with prolonged exposure. For one tested batch, the initial purity of 97.7% showed minimal change after one week at 40 °C/75% RH (96.8% open, 97.3% closed), while a significant decrease was observed at 60 °C (58.4%). Under accelerated conditions (40 °C/75% RH, open and closed, and 60 °C) over two weeks, the crystal form did not change (evidenced by PXRD). As anticipated, pH dependent aqueous and biorelevant fluid (FesSIF, FasSIF, SGF) solubility is fairly high (≥ 2 mg/mL) for mCMX110; enabling up to 100 mg/mL homogeneous suspensions in simple aqueous media (f.ex, 0.5% MC or 0.5% NaCMC) that worked well in PK studies (**Figure 3, Table 2**). Regarding stability in biorelevant media, **mCMX110** was very stable in SIF and SGF, whereas PF-00835231 was found (**Table 2**) to have poor SIF stability, which may in part contribute to the poor oral bioavailability of this comparator.

**Table 2:**
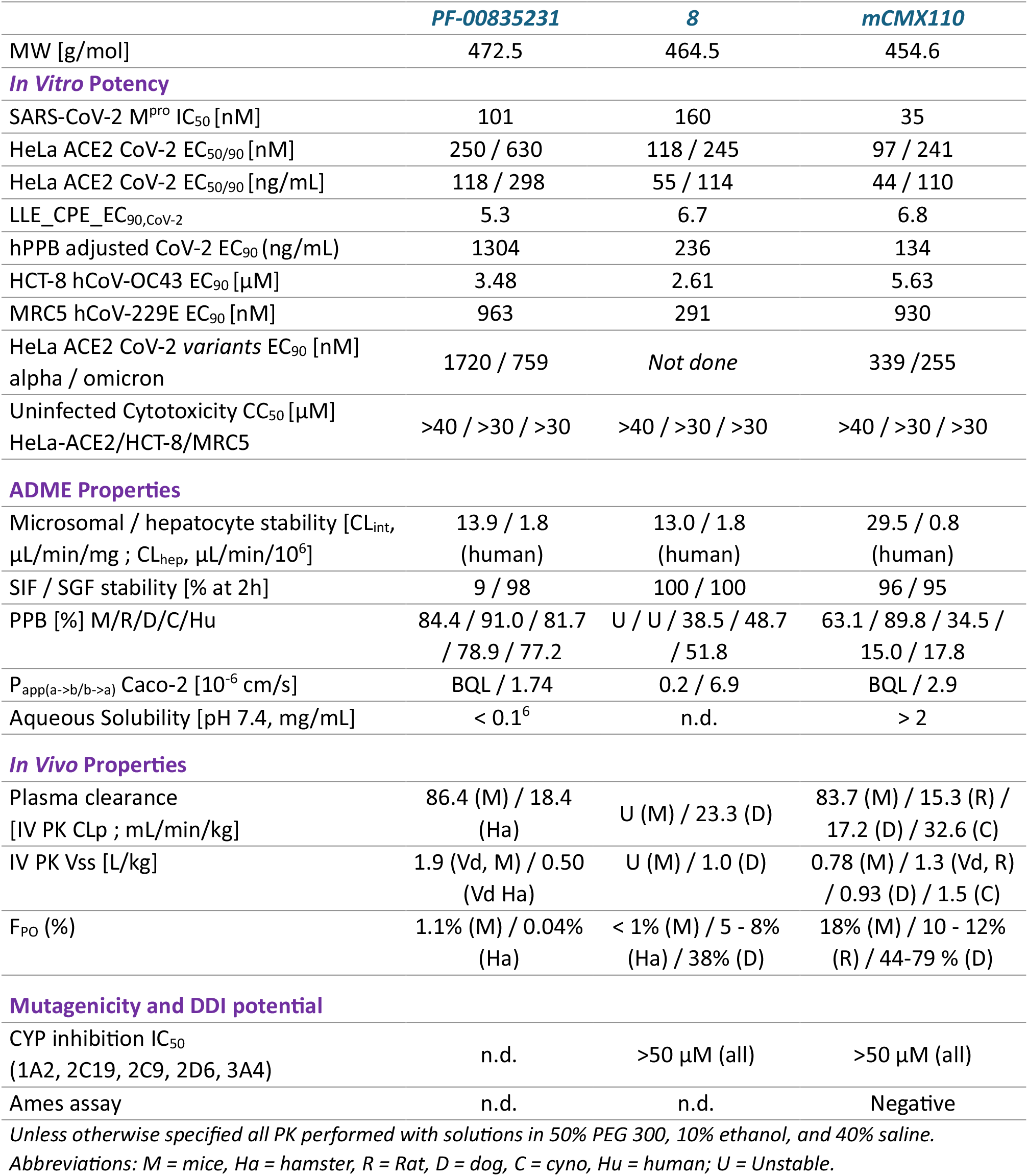
Comparative performance of HMK compounds 8 and mCMX110 vs. PF-00835231.

|  | *PF-00835231* | *8* | *mCMX110* |
| --- | --- | --- | --- |
| MW [g/mol] | 472.5 | 464.5 | 454.6 |
| ***In Vitro Potency*** |  |  |  |
| SARS-CoV-2 M<sup>pro</sup> IC<sub>50</sub> [nM] | 101 | 160 | 35 |
| HeLa ACE2 CoV-2 EC<sub>50/90</sub> [nM] | 250 / 630 | 118 / 245 | 97 / 241 |
| HeLa ACE2 CoV-2 EC<sub>50/90</sub> [ng/mL] | 118 / 298 | 55 / 114 | 44 / 110 |
| LLE_CPE_EC<sub>90,CoV-2</sub> | 5.3 | 6.7 | 6.8 |
| hPPB adjusted CoV-2 EC<sub>90</sub> (ng/mL) | 1304 | 236 | 134 |
| HCT-8 hCoV-OC43 EC<sub>90</sub> [μM] | 3.48 | 2.61 | 5.63 |
| MRC5 hCoV-229E EC<sub>90</sub> [nM] | 963 | 291 | 930 |
| HeLa ACE2 CoV-2 *variants* EC<sub>90</sub> [nM]<br>alpha / omicron | 1720 / 759 | *Not done* | 339 / 255 |
| Uninfected Cytotoxicity CC<sub>50</sub> [μM]<br>HeLa-ACE2/HCT-8/MRC5 | >40 / >30 / >30 | >40 / >30 / >30 | >40 / >30 / >30 |
| ***ADME Properties*** |  |  |  |
| Microsomal / hepatocyte stability [CL<sub>int</sub>,<br>μL/min/mg ; CL<sub>hep</sub>, μL/min/10<sup>6</sup>] | 13.9 / 1.8<br>(human) | 13.0 / 1.8<br>(human) | 29.5 / 0.8<br>(human) |
| SIF / SGF stability [% at 2h] | 9 / 98 | 100 / 100 | 96 / 95 |
| PPB [%] M/R/D/C/Hu | 84.4 / 91.0 / 81.7<br>/ 78.9 / 77.2 | U / U / 38.5 / 48.7<br>/ 51.8 | 63.1 / 89.8 / 34.5 /<br>15.0 / 17.8 |
| P<sub>app(a->b/b->a)</sub> Caco-2 [10<sup>-6</sup> cm/s] | BQL / 1.74 | 0.2 / 6.9 | BQL / 2.9 |
| Aqueous Solubility [pH 7.4, mg/mL] | < 0.1<sup>6</sup> | n.d. | > 2 |
| ***In Vivo Properties*** |  |  |  |
| Plasma clearance<br>[IV PK CL<sub>p</sub> ; mL/min/kg] | 86.4 (M) / 18.4<br>(Ha) | U (M) / 23.3 (D) | 83.7 (M) / 15.3 (R) /<br>17.2 (D) / 32.6 (C) |
| IV PK V<sub>ss</sub> [L/kg] | 1.9 (Vd, M) / 0.50<br>(Vd Ha) | U (M) / 1.0 (D) | 0.78 (M) / 1.3 (Vd, R)<br>/ 0.93 (D) / 1.5 (C) |
| F<sub>PO</sub> (%) | 1.1% (M) / 0.04%<br>(Ha) | < 1% (M) / 5 - 8%<br>(Ha) / 38% (D) | 18% (M) / 10 - 12%<br>(R) / 44-79 % (D) |
| ***Mutagenicity and DDI potential*** |  |  |  |
| CYP inhibition IC<sub>50</sub><br>(1A2, 2C19, 2C9, 2D6, 3A4) | n.d. | >50 μM (all) | >50 μM (all) |
| Ames assay | n.d. | n.d. | Negative |

*Unless otherwise specified all PK performed with solutions in 50% PEG 300, 10% ethanol, and 40% saline.*
*Abbreviations: M = mice, Ha = hamster, R = Rat, D = dog, C = cyno, Hu = human; U = Unstable.*

**Figure 3:**
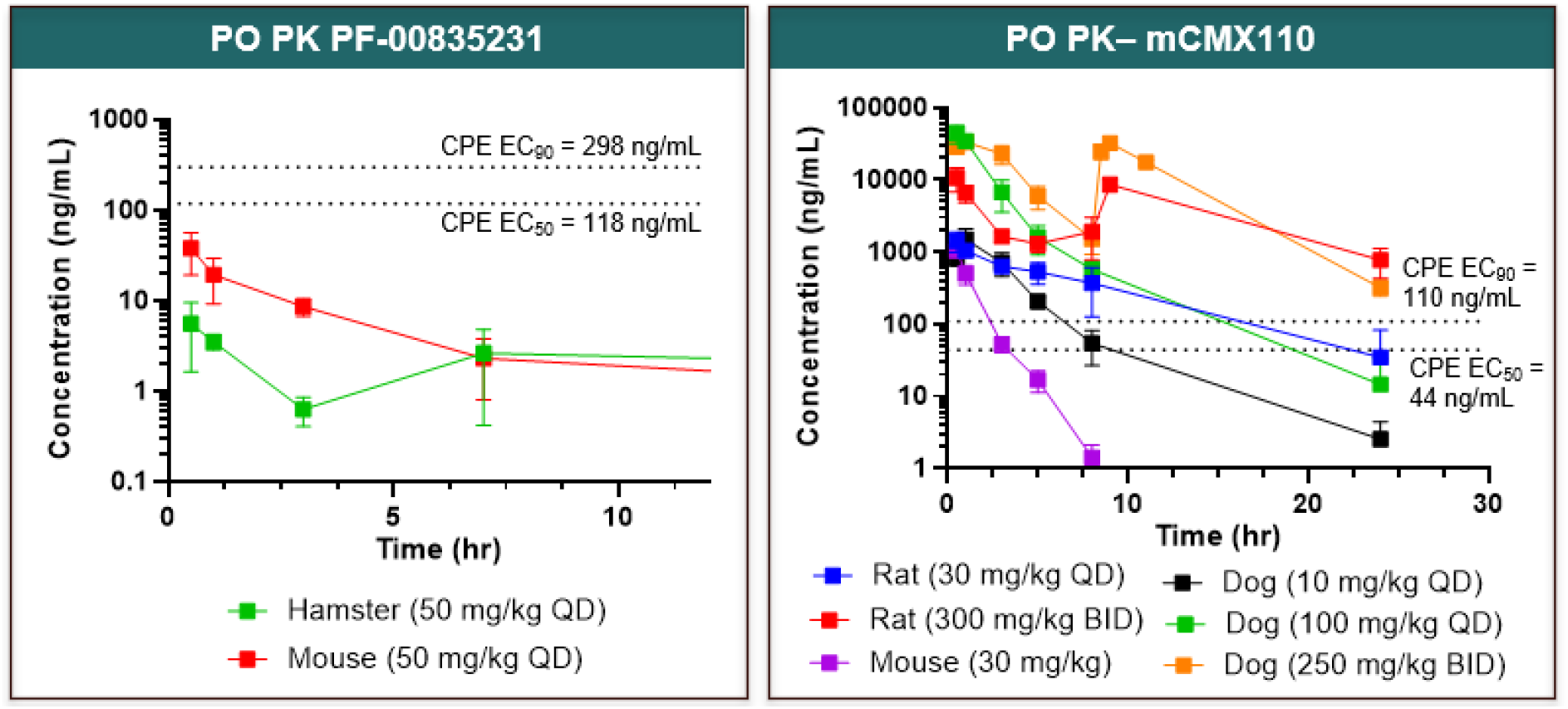
Oral preclinical pharmacokinetic profile of mCMX110 is superior to PF-00835231. **mCMX110** shows superior coverage over EC_90_ and has good dose-proportionality to exposure in both rats and dogs. Except for otherwise specified, all represented PK studies done in 50% PEG300, 10% Ethanol, and 40% Saline. mCMX110 dog 100 mg/kg QD and 250 mg/kg BID oral dosing done using suspension formulations in 0.5% NaCMC (aqueous). **mCMX110** rat 300 mg/kg BID oral dosing done using suspension formulations in 0.5% MC (aqueous). Oral bioavailability (F_PO_): PF-00835231 = 0.04% (hamster), 1.1% (mouse); **mCMX110** = 18% (mouse), 10 – 12% (rat), 44 – 79% (dog).

### Preclinical oral pharmacokinetics of mCMX110 outperforms PF-00835231

Dosed orally in simple aqueous suspensions, **mCMX110** readily covers its SARS-CoV-2 EC_90_ threshold target (**Figure 3**). In mice, as anticipated, faster clearance makes the coverage less profound in low dose (30 mg/kg). Still, even in mice the plasma exposure above the EC_90_ threshold for **mCMX110** is improved relative to **PF-00835231**. In addition, **mCMX110** showed good dose-escalation feasibility in both rats and dogs, resulting in good proportionality to exposure increase. In dog PO PK, **mCMX110** demonstrated consistently high oral bioavailability (44-79%) across multiple studies. Notably, early aryl oxamide lead **8** had better performance than **PF-00835231** (supporting information); but generally lacked stability in rodent plasma (**Table 2**). This was another key factor to supporting the selection of mCMX110 as the lead compound, where the plasma stability issue was solved.

### mCMX110 has a good preclinical safety profile *in vitro* and *in vivo*

In vitro potency profile of **mCMX110** was not driven by any cytotoxicity, where CC_50_ was always above the top concentrations measured in representative assays (> 30 µM, **Table 2**). Oral administration of **mCMX110** in dogs at 125 mg/kg/dose for 5 days BID did not result in adverse effects including on behavioral, hematologic, renal, or coagulation parameters. Similarly, repeat-dose tolerability study in rats for 5 days (100 – 300 mg/kg/dose BID PO) did not reveal any prohibitive findings. **mCMX110** had no effect on inhibition of five Cyp isoforms tested and was negative for mutagenicity in a mini-Ames assay (**Table 2**).

## 4. Discussion

Conventionally, hydroxymethyl ketone (HMK) warhead bearing compounds are often perceived as less favorable for oral delivery, given the presence of additional hydrogen bond donor in the molecule, and the ensuing polarity and potential metabolic liabilities. The present work expands the chemical space of SARS-CoV-2 Mpro inhibitors by revisiting this HMK warhead space and demonstrating that their historical limitations can be mitigated through holistic optimization of molecular properties rather than warhead substitution alone. While recent clinical success has largely centered on nitrile-based inhibitors, our findings highlight that HMKs remain a viable and differentiated modality with favorable selectivity and tunable pharmacokinetic properties.

In this work, **mCMX110** was identified as a lead compound that achieves a balanced profile across potency, physicochemical properties, and *in vivo* exposure. Compared to **PF-00835231, mCMX110** demonstrates ∼3x improved antiviral potency both biochemically and in cells. Importantly, these gains were achieved without increasing lipophilicity, as reflected in maintained or improved ligand efficiency metrics. In addition, **mCMX110** also has reduced plasma protein binding, which helps drive a substantially higher unbound fraction *in vivo*. This improvement in free drug exposure is particularly relevant in the context of antiviral efficacy, where target engagement is governed by unbound concentrations. Combined, the greater unbound exposure/potency may translate into improved antiviral efficacy for this compound in future studies.

Our SAR exploration underscored some key design principles for HMK warhead based Mpro inhibitors. This is important as our extensive work in this area has established that not all warheads pair similarly with different P2/P3 groups. Aryl oxamide motifs as P2-caps provided marked potency gains, consistent with potential favorable interactions within the S3/S4 pockets,^11^ resulting in early lead **8**. However, these aryl oxamides drove mutagenicity signals in Ames assays, and had general plasma instability in rodents. The eventual transition to non-anilinic oxamides in **mCMX110** preserved potency while resolving these liabilities, illustrating the importance of balancing target engagement with developability constraints. Expansion of the P1 lactam from a five-to six-membered ring contributed to potency improvements, likely by better complementing the S1 subsite geometry. Further expansion to a seven-membered ring (e.g., **21**) resulted in loss of potency.

A distinguishing feature of **mCMX110** is its favorable oral pharmacokinetic profile, achieved without the need for prodrug strategies or pharmacokinetic boosters. Unlike **PF-00835231**, which suffers from poor oral exposure and instability in simulated intestinal fluid; **mCMX110** exhibits good stability in simulated intestinal and gastric fluids and has oral exposure reaching >50% in higher animals. mCMX110 also has excellent aqueous solubility (> 2 mg/mL) and suspendability, enabling robust formulability as simple aqueous suspensions. These properties subsequently translate into robust *in vivo* exposure across species, with oral dosing achieving plasma concentrations exceeding the antiviral EC_90_ threshold. Notably, dose-proportional exposure and sustained target coverage were observed in both rodent and non-rodent species, supporting the potential for clinically relevant oral dosing regimens.

From a safety perspective, **mCMX110** demonstrated a clean *in vitro* profile, with no observable cytotoxicity at concentrations well above antiviral potency and no detectable CYP inhibition across major isoforms. Furthermore, repeat oral dosing in rats and dogs (5 days BID) was well tolerated, with no adverse findings in clinical chemistry or behavioral assessments. Although additional toxicological evaluation is required, these early results are encouraging and consistent with the generally favorable selectivity profile of HMK warheads.

It is our view that further optimization of this scaffold is likely feasible to improve upon potency and stability properties of **mCMX110**. Ultimately, we intend to put this compound (or an improved analog thereof) in a preclinical efficacy study to establish proof-of-concept efficacy data. The improved analogs will include Lufotrelvir-like prodrug modifications that may enhance the profile of **mCMX110** even further.

## 5. Conclusion

This work demonstrates that hydroxymethyl ketone warhead-based compounds can be advanced as orally bioavailable antivirals with competitive potency and favorable pharmacological properties. The discovery of **mCMX110** provides a compelling example of how revisiting underexplored warheads can yield differentiated candidates that address limitations of current therapies. More broadly, these findings reinforce the importance of maintaining chemical diversity in antiviral drug discovery to ensure preparedness against current and future coronavirus threats.

## Supplementary Materials

The supporting information can be downloaded online

## Author Contributions

Conceptualization, N.E., K.W, G.G., A.G., A.R., C.M., J.C., A.C.; methodology, N.E., K.W., L.R., J.C., A.C.; validation, N.E., K.W.; formal analysis, N.E., K.W., S. J., K.L., S.L.; investigation, N.E., K.W., F.W., S.G., G.G., K.W., L.R., A.W., J.P., A.N., Y.L., W.M., L.S., N.O., J.M., M.B., M.K., A.G., E.H., V.N., V.C., A.R.; writing – original draft preparation, N.E.; writing – review and editing, N.E., K.W., K.L.; visualization, N.E.; supervision, N.E., K.W., J.C., A.C.; funding acquisition, A.C., P.S.; project administration, A.C.. All authors have read and agreed to the published version of the manuscript.

## Acknowledgments

This work was supported by grants from the Bill and Melinda Gates Foundation (INV-028691, COVID-19 CTA GPP). The authors would like to thank the diverse team of subject-matter-experts engaged by the Bill and Melinda Gates Foundation, including senior program officer Dr. Peter Warner. In addition, we’d also like to thank rest of the Calibr-Skaggs colleagues who supported our protease inhibitor program, particularly Dr. Kelli L. Kuhen and Dr. Eugenio De Hostos.

## Abbreviations

The following abbreviations are used in this manuscript:

ADME: Absorption, Distribution, Metabolism, and Excretion
CPE: Cytopathic effect
DMPK: Drug Metabolism and Pharmacokinetics
DSC: Differential Scanning Calorimetry
FesSIF: Fed-state simulated intestinal fluid
FasSIF: Fasted-state simulated intestinal fluid
HLM: Human liver microsomes
HMK: Hydroxymethyl ketone
IV: Intravenous
LLE: Lipophilic ligand efficiency
MC: Methylcellulose
Mpro: Main protease
NaCMC: Sodium carboxymethyl cellulose
NMR: Nuclear magnetic resonance
PK: Pharmacokinetics
PO: Oral administration
PPB: Plasma protein binding
RH: Relative humidity
SAR: Structure-Activity Relationship
SARS-CoV: Severe acute respiratory syndrome coronavirus
SFC: Supercritical fluid chromatography
SGF: Simulated gastric fluid
SIF: Simulated intestinal fluid
Vd: Volume of distribution
Vss: Volume of distribution at steady state
WH: Warhead

