## supporting information for "Orally Bioavailable SARS-CoV-2 Protease Inhibitors Bearing a Hydroxymethyl Ketone Warhead"

**for**

### mCMX110: Spectral Data

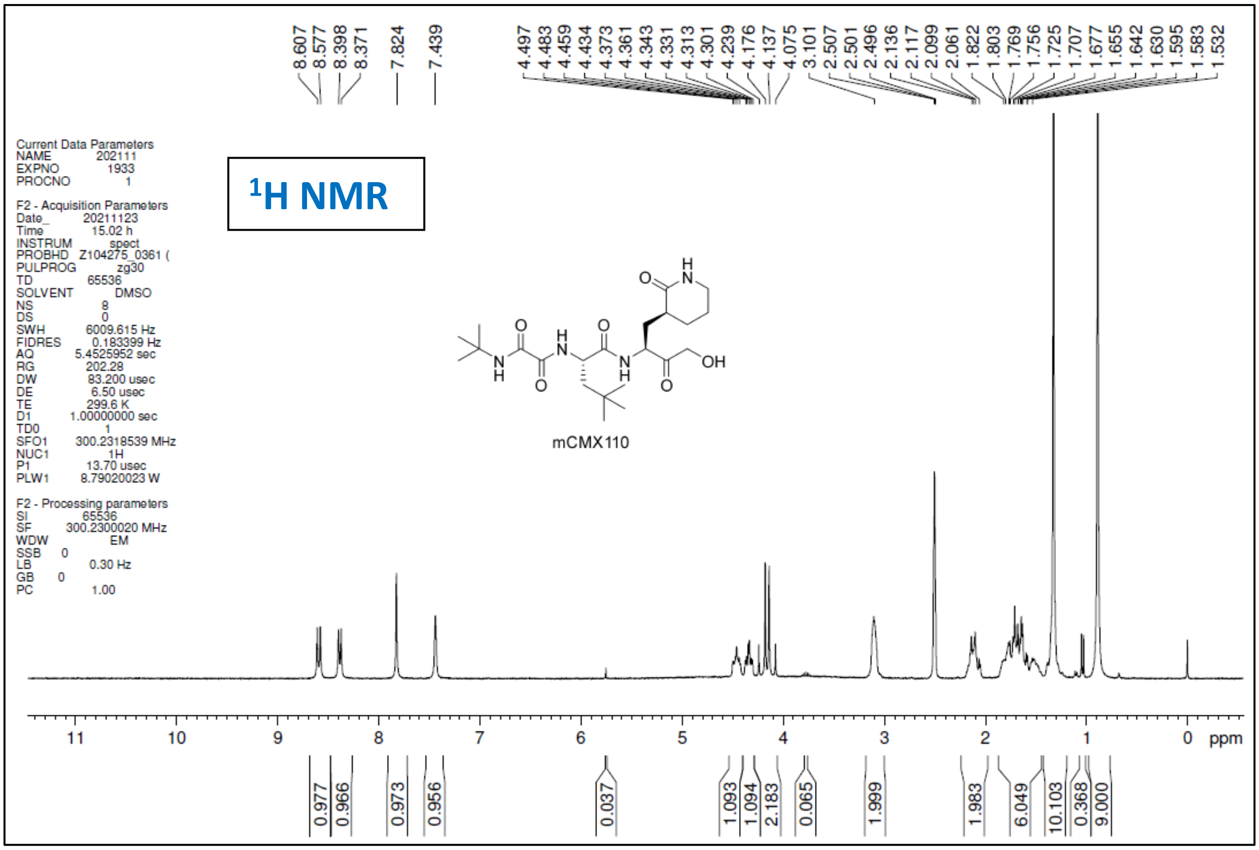


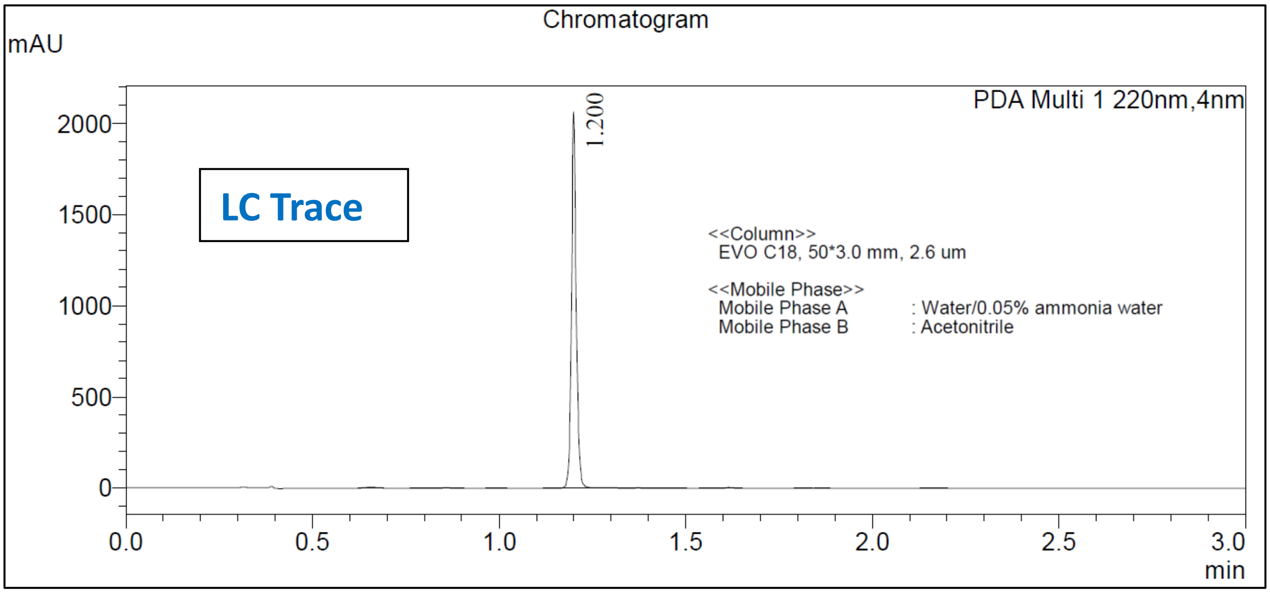

Mass Spectrum
Line#: 1
Acquisition Mode: Scan (Positive)
R.Time: 1.200(Scan#:185) Spectrum Mode: Averaged 1.193-1.207(184-186)
BasePeak: 455.3(451362)

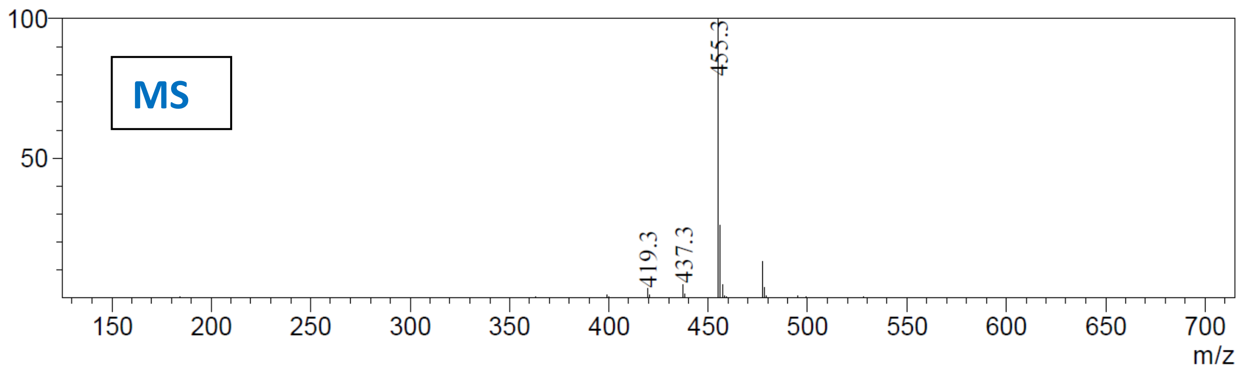


#### SFC-MS Analysis

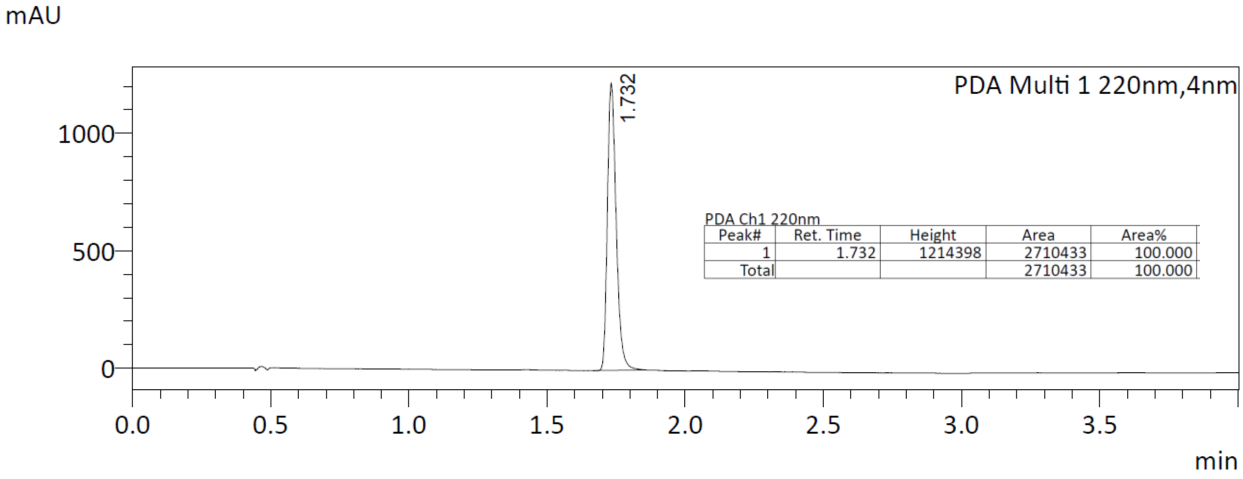


PDA Ch1 220nm

| Peak# | Ret. Time | Height | Area | Area% |
|---|---|---|---|---|
| 1 | 1.732 | 1214398 | 2710433 | 100.000 |
| Total | | | 2710433 | 100.000 |

MS Spectrum
Line#:1 R.Time:1.730(Scan#:263)
MassPeaks:288
Spectrum Mode:Single 1.730(263) Base Peak:455.10(20055)
BG Mode:None Segment 1 - Event 1

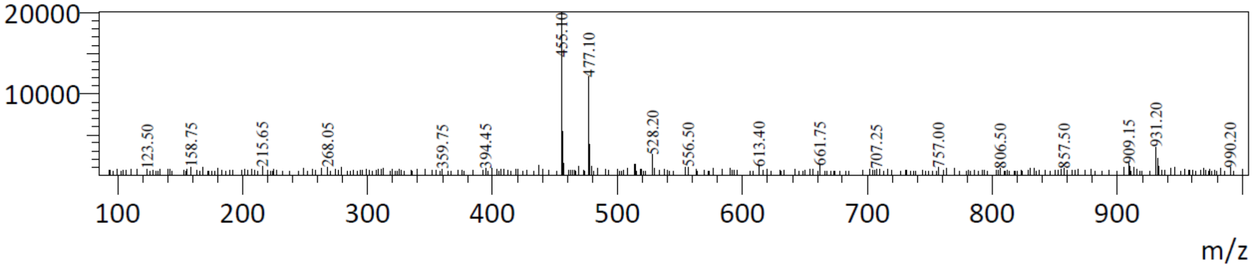

#### Oral PK profile of 8 in Hamster and Dog

- Compound was unstable in mouse and rodent plasma, necessitating evaluation in hamsters to obtain measurable PK exposure in rodents.
- Dosing formulation for all studies = 50% PEG 300, 10% ethanol, and 40% saline.
- Dose Normalized IV AUC = 158 hr.ng/mL; Hamster $F_{PO}$ = 5-8%; Dog $F_{PO}$ = 38%

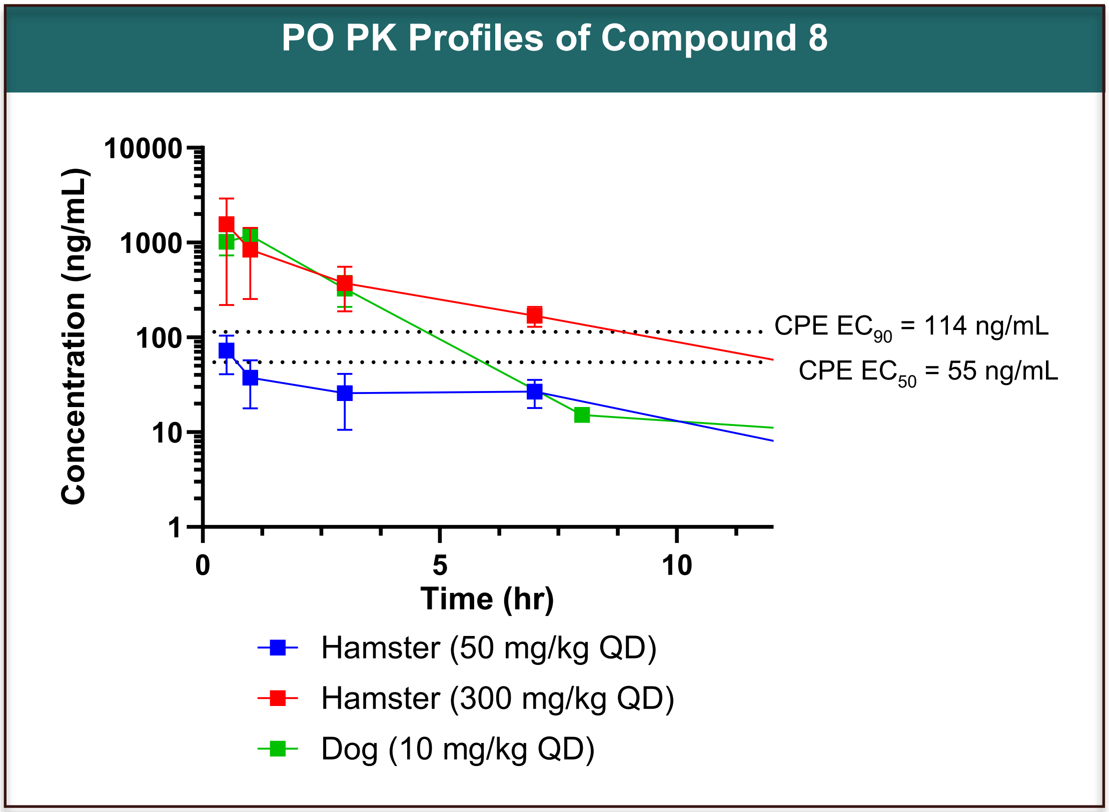
